# scTopoGAN: unsupervised manifold alignment of single-cell data

**DOI:** 10.1101/2022.04.27.489829

**Authors:** Akash Singh, Marcel J.T. Reinders, Ahmed Mahfouz, Tamim Abdelaal

**Affiliations:** Delft Bioinformatics Lab, Delft University of Technology, 2628 XE Delft, The Netherlands; Leiden Computational Biology Center, Leiden University Medical Center, 2333ZC Leiden, The Netherlands; Department of Human Genetics, Leiden University Medical Center, 2333ZC Leiden, The Netherlands; LKEB, Department of Radiology, Leiden University Medical Center, 2333ZC Leiden, The Netherlands

## Abstract

**Motivation:** Single-cell technologies allow deep characterization of different molecular aspects of cells. Integrating these modalities provides a comprehensive view of cellular identity. Current integration methods rely on overlapping features or cells to link datasets measuring different modalities, limiting their application to experiments where different molecular layers are profiled in different subsets of cells.

**Results:** We present scTopoGAN, a method for unsupervised manifold alignment of single-cell datasets with non-overlapping cells or features. We use topological autoencoders to obtain latent representations of each modality separately. A topology-guided Generative Adversarial Network then aligns these latent representations into a common space. We show that scTopoGAN outperforms state-of-the-art manifold alignment methods in complete unsupervised settings. Interestingly, the topological autoencoder for individual modalities also showed better performance in preserving the original structure of the data in the low-dimensional representations when compared to other manifold projection methods. Taken together, we show that the concept of topology preservation might be a powerful tool to align multiple single modality datasets, unleashing the potential of multi-omic interpretations of cells.

**Availability and implementation:** Implementation available on GitHub (https://github.com/AkashCiel/scTopoGAN). All datasets used in this study are publicly available.

## 1 Introduction

A growing number of single-cell technologies allow the characterization of distinct molecular features of cells, such as single-cell RNA-sequencing (scRNA-seq) or measuring chromatin accessibility at single-cell resolution (scATAC-seq). Despite advances in multimodal technologies (Zhu *et al*., 2020), these molecular features are mostly measured from different subsets of cells. Sometimes the measured modalities share common features, for example when spatial transcriptomics and scRNA-seq are applied on the same tissue. Because the datasets are not measured from the same cells, they have to be aligned into a common space using data integration methods (Argelaguet *et al*., 2021).

Multi-omic data integration methods aim to find a joint latent space representing information from multiple modalities. These methods include MOFA+ (Argelaguet *et al*., 2020), Seurat WNN (weighted nearest neighbor) (Hao *et al*., 2021), totalVI (Gayoso *et al*., 2021) and Mixture of Experts (Shi *et al*., 2019). These methods require the cell-cell correspondence between the different omics modalities and cannot be applied when multiple unimodal assays are used to profile different cells from the same biological sample. This problem is referred to as diagonal integration, and represents the most challenging case of single-cell multi-omics data integration (Argelaguet *et al*., 2021).

Previous methods, such as MATCHER (Welch *et al*., 2017), SCIM (Stark et al., 2020), UnionCom (Cao et al., 2020) and MMD-MA (Singh et al., 2020), have addressed this challenging integration task by assuming a similar cellular composition between unimodal datasets collected from the same tissue. MATCHER uses Gaussian processes to embed cells from multiple modalities onto a 1D trajectory. SCIM uses variational autoencoders with an adversarial objective function to learn a modality-invariant latent representation. UnionCom first defines geometrical matches between cells across different modalities and projects the different features onto a common latent representation which is comparable for the matched cells. MMD-MA minimizes the maximum mean discrepancy between the different modalities in the learned latent space. These multi-modal alignment methods, however, suffer from several limitations. MATCHER and SCIM are not fully unsupervised as they require (partial) cell type annotations for each of the different modalities in order to align the data. Additionally, MATCHER can only align 1D trajectory structures and cannot deal with more complex structures. UnionCom and MMD-MA represent fully unsupervised manifold alignment methods, however, both methods were tested on single-cell multi-omics data, in which the multiple modalities were measured from the same cell, with perfect cell-to-cell correspondences. Although they did not exploit this correspondence in their methods, the integration performance drops significantly (as we show later) when, more realistic, datasets lacking this correspondence are used. Recently, a new category of diagonal integration methods emerged, including bridge integration (Hao *et al*., 2022), UINMF (Kriebel and Welch, 2022) and StabMap (Ghazanfar *et al*., 2022). These methods require an additional single-cell multi-modal dataset containing cell-cell correspondence and measuring the same modalities of the unimodal datasets to be diagonally integrated. This additional multi-modal dataset can act like a bridge and translate information between the distinct unimodal datasets guiding the data integration process. Moreover, GLUE (Cao and Gao, 2022) is using a graph-based modeling of the regulatory interactions in order to integrate gene expression and chromatin accessibility data. These methods are beyond the scope of our study since they require an additional representative multi-modal layer.

Considering different single-cell modalities measured from the same biological sample, the main assumption in integration is that the different modalities lie on the same underlying manifold (Sun *et al*., 2018). Preserving the topology of the datasets is crucial when constructing and integrating the different manifolds. Since the different modalities are measuring distinct features, it is necessary to first find a low-dimensional representation of each modality separately. Topological autoencoders have been recently introduced to project high-dimensional data into a low-dimensional latent space while preserving the data topology (Moor *et al*., 2020). Next, these low-dimensional manifolds have to be aligned into a common space with minimal distortion to the original topology of each data modality. Generative Adversarial Networks (GANs) were successfully used in the computer vision field (Gui *et al*., 2021). GANs were previously used to project biological datasets onto each other (Amodio and Krishnaswamy, 2018), however, based on correspondence information between the datasets, and not in a fully unsupervised setting.

We propose scTopoGAN, a topology-preserving multi-modal alignment of two single-cell modalities with non-overlapping cells or features. scTopoGAN first finds topology-preserving latent representations of the different modalities, which are then aligned in an unsupervised way using a topology-guided GAN. scTopoGAN is fully unsupervised with no requirement for cell type annotations. Our results show that scTopoGAN outperforms state-of-the-art methods, producing joint representation of distinct datasets with better matching between cellular populations.

## 2 Methods

### 2.1 scTopoGAN overview

scTopoGAN is designed to align two datasets measuring two different single-cell modalities, each measured on different non-matching cells. scTopoGAN consists of two steps: 1) manifold projection, and 2) manifold alignment (Fig. 1). Assuming a lower-dimensional manifold structure for single-cell datasets (Bac and Zinovyev, 2019), scTopoGAN first finds the manifold for each modality separately, with explicit preservation of the data topology. Then, the latent space representation of the two modalities are aligned in a topology-preserving manner, exploiting the assumption that the topology of the cells in the two modalities is the same. This alignment step should preserve relevant inter-modality correspondence such that similar cell types should be aligned between different modalities.

**Fig. 1.**
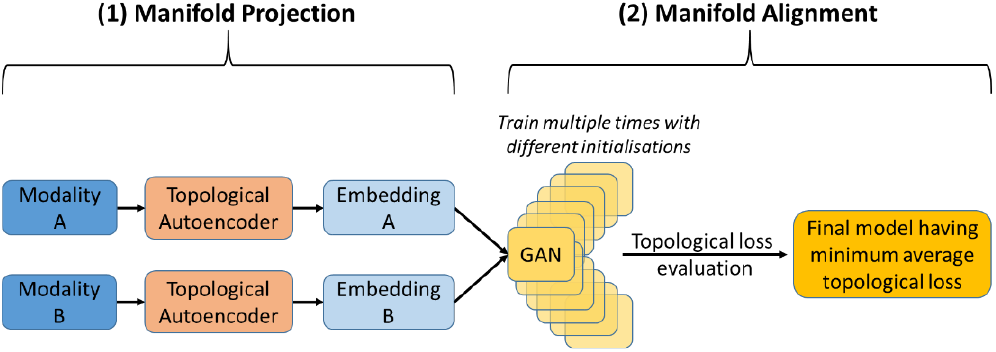
scTopoGAN overview. **sc**TopoGAN consists of two stages: (1) Manifold projection using Topological autoencoders to obtain a low dimensional embedding (manifold) for each modality independently. (2) Manifold alignment using a GAN. Hereto, 20 different GAN models are trained with random initializations. Then that model is selected which has the minimum average topological loss. The selected model is further trained for 1000 additional epochs to produce the final alignment.

#### 2.1.1 Manifold projection

To project each modality to a lower-dimensional latent space, scTopoGAN uses a topological autoencoder (topoAE) (Moor *et al*., 2020), which chooses point-pairs that are crucial in defining the topology of the manifold instead of trying to optimize all possible point-pairs. A topoAE is based on the concept of persistence homology (Edelsbrunner and Harer, 2008) which selectively considers edges connecting point-pairs below a certain distance threshold. These edges are used to construct local neighborhoods together constituting large-scale topological features. By repeating this procedure for a wide range of distance thresholds, persistent topological features are defined, where the point-pairs constituting them are known as persistence pairings. Preserving the distances between these pairings in a lower-dimensional projection of the data preserves the data topology. The loss function of the topoAE is defined as:

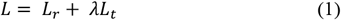

where *L*_*r*_ is the reconstruction loss between the input and reconstructed output of the autoencoder across all cells, and *L*_*t*_ represents the topological loss, with *λ* is the weight of the topological loss. The topological loss is defined as:

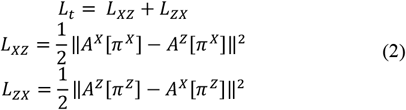

where *X* is the original input data and *Z* is the encoded latent representation, *A*^*X*^ and *A*^*Z*^ are the distance matrices in the original and latent spaces respectively, *π* ^*X*^ and *π*^*Z*^ are the persistence pairings in the original and latent spaces respectively. *A*·[*π*·] represent subset of distances in the space *A*· defined by the topologically relevant edges in that space *π*·. The term *L*_*XZ*_ ensures that persistence pairings relevant to the original manifold are equidistant in both the original and the latent spaces, while *L*_*ZX*_ ensures that persistence pairings relevant to the latent manifold are equidistant in both spaces.

#### 2.1.2 Manifold alignment

We used a GAN (Goodfellow *et al*., 2014) to align one modality (source) to the other modality (target). The generator part of the GAN aims to project the source modality onto the target modality, resulting in a combined dataset. We use a single hidden layer generator network against a double hidden-layer discriminator network. The GAN was trained for 1000 epochs.

To ensure a topology-preserving alignment of the two modalities, we trained 20 different GANs and selected the GAN which best preserves the topological loss (Eq. 2) between the source data and its projection in the target data space. To do so, for each GAN, the topological loss is calculated from epoch 500 every 100^th^ epoch until epoch 1000 (6 values) and then averaged. The topological loss was calculated for a batch size of 1000 to balance between coverage of global structure in each batch and compute memory requirements. The generator network of the selected GAN is then loaded into a new GAN model with a new discriminator network as its adversary. This final model is then trained for an additional 1000 epochs to obtain the final aligned manifolds.

### 2.2 Datasets

#### 2.2.1 Peripheral blood mononuclear cells (PBMC) dataset

The PBMC dataset consists of healthy human PBMCs, simultaneously profiling gene expression (RNA) and chromatin accessibility (ATAC) from the same cells using the 10x multiome protocol. The dataset was downloaded from the 10x Genomics website (https://support.10xgenomics.com/single-cell-multiome-atac-gex/datasets/1.0.0/pbmc_granulocyte_sorted_10k). The **Full PBMC** dataset contained 11,910 cells, profiling 36,601 genes and 108,377 peaks, having one-to-one correspondence between the two modalities, and including 7 major cell classes (CD4 T cells, CD8 T cells, Monocytes, NK cells, Dendritic cells, B cells and HSPC) which are further divided into 20 cell subclasses.

To simulate a realistic data in which the cell-cell correspondences do not exist between the two modalities, we generated the **Partial PBMC** dataset where we randomly removed 30% of the cells (after preprocessing) from both RNA and ATAC independently stratified across the different cell classes. This results in a total of 7,349 cells for each modality, including 2,100 cells which have no corresponding cells in the other modality. In this case, the **Partial PBMC** dataset represents an example with partial correspondence.

#### 2.2.2 Bone marrow (BM) dataset

The original BM dataset consists of human bone marrow cells, simultaneously profiling gene expression (RNA) and protein expression (antibody-derived tags, ADT) from the same cells using the CITE-seq protocol (Stoeckius *et al*., 2017). The dataset contained 30,672 cells, profiling 17,009 genes and 25 ADT, with cell-cell correspondence between both modalities, and including 5 major cell classes (T cells, B cells, Mono/DC cells, NK cells and Progenitor cells), further categorized into 27 different cell subclasses. For the purpose of our study, we randomly selected 10,235 cells from each modality independently in a stratified manner across the cell classes. In this case, the **BM** dataset does not contain any cell-cell correspondences between the two modalities.

#### 2.3 Data preprocessing

We performed all data preprocessing using the Seurat v4.0 R package (Hao *et al*., 2021). For the PBMC dataset, we filtered out cells with RNA count below 1,000 or above 25,000, cells with ATAC count below 5,000 or above 70,000, and cells with mitochondrial percentage above 20%, resulting in a total of 10,412 cells. Further, the RNA modality is normalized using `SCTransform` (Hafemeister and Satija, 2019), selecting the top 3000 variable genes. The ATAC modality was normalized using the `RunTFIDF` function using a scaling factor of 10,000, followed by finding the top peaks using the `FindTopFeatures` function with min.cutoff = q0. Next, we reduced the dimensionality of the RNA and ATAC data to 50 dimensions using Principal Component Analysis (PCA) and Latent Semantic Indexing (LSI), respectively. These 50-dimensional datasets are used as input to the scTopoGAN workflow.

For the BM dataset, the RNA modality was normalized using a scaling factor of 10,000 followed by log-transformation. The top 2000 variable genes were selected, next the data was scaled and centered. The ADT modality was centered log-ratio (CLR) normalized, scaled and centered. The RNA data was reduced to 50 dimensions using PCA, while dimensionality reduction was not necessary for the ADT modality which only had 25 features.

### 2.4 Benchmarking methods

For the manifold projection step, we compared the performance of the topoAE with a standard variational autoencoder (VAE) (Kingma and Welling, 2014), which is used for manifold projection in SCIM (Stark *et al*., 2020), and a regular autoencoder (AE). Further, we used UMAP (McInnes *et al*., 2018) as a base-line for the manifold projection evaluation. Next, we compared the alignment performance of scTopoGAN with the state-of-the-art methods UnionCom and MMD-MA.

### 2.5 Evaluation metrics

To evaluated the manifold projection, we used the Silhouette score (Rousseeuw, 1987) which assesses the separation between the cell classes. The Silhouette score ranges from -1 to 1, where a higher value indicates better separated classes. Additionally, we calculated the Kullback-Leibler divergence *KL*_*σ*_ between the density estimates of the input data and its latent space representation (Moor *et al*., 2020). The *KL*_*σ*_ calculation requires the pairwise distance matrices of the original input data and its latent representation. Gaussian kernel of size *σ* (we used *σ* = 0.01) is applied for each point to estimate its density based on the distances to other points. The *KL*_*σ*_ value quantifies the dissimilarity between the density estimates in both spaces (input and latent), thus lower values (≈ 0) indicate better manifold projection performance.

To evaluate the manifold alignment, for each cell in one modality, we determine its k-neighboring cells from the other modality in the final aligned common space (k=5, Euclidean distance). Next, we compare the class/subclass annotation of that cell with the majority vote of its neighbors and check whether it is a match or not. We report the percentage of cells with matching cell class/subclass denoted as the Celltype matching and the Subcelltype matching scores, respectively.

### 2.6 Implementation details

To train the topoAE, we used a learning rate of 1e-03, batch size of 50, latent size of 8 dimensions, and an architecture of two hidden layers followed by batch normalization and ReLU activation. For the hyperparameter *λ*, we tested values ranging from 0.5 to 3.0 as recommended by (Moor et al., 2020). The hidden layers are both of size 32, except for the **BM** ADT data (input dimensions = 25), the size of the hidden layers is 16. We used the same architecture for VAE and AE. For all autoencoder models, we split the data into 80% training and 20% validation. We trained the models for a minimum of 50 epochs and a maximum of 200 epochs, with an early stop if the validation loss did not improve for 10 consecutive epochs (after the initial 50 epochs).

For the GAN model, we used a generator hidden layer of size 30 and a discriminator hidden layers of sizes 60 and 30, batch size of 512 and learning rates of 1e-03 and 1e-02 for the generator and discriminator, respectively. We followed previous work using GANs to stabilize the training process (Radford *et al*., 2016) by sampling initialization weights from a normal distribution N(0,0.02), and using a Leaky ReLU as the activation function for the discriminator with an activation value of 0.2, while using ReLU activation for the generator. UnionCom and MMD-MA were trained for 1000 epochs using default hyperparameters settings. For all models, input datasets were randomly shuffled to ensure cell-cell correspondence is not implicitly provided to the model.

## 3 Results

### 3.1 Topological autoencoder produced better manifold projections compared to other methods

Before integrating different data modalities, it is crucial to acquire a proper low-dimensional embedding of each modality separately. For this manifold projection task, we used a topoAE which has been shown to produce reliable topology approximations (Moor *et al*., 2020). To the best of our knowledge, topoAEs have not been applied on biological datasets which, compared to classical datasets used in machine learning, contain continuous topological structures (Rizvi *et al*., 2017). Using both RNA and ATAC modalities of the **Full PBMC** dataset, and both RNA and ADT of the **BM** dataset, we compared the manifold projection performance of the topoAE with three other methods (Table 1). All methods were used to reduce the 50-dimensional (PCA or LSI) data or the 25-dimensional ADT data to 8 dimensions, additionally UMAP was used to produce 2-dimensional embedding for visualization purpose. Furthermore, we tested different settings for the topological loss weight *λ* of the topoAE. Results show that topoAE is the best method in preserving the original data density estimates having overall the lowest *KL*_0·01_ value, except for the PBMC ATAC data, where VAE is performing better in terms of density preservation. However, UMAP obtained the highest Silhouette score producing better separation between different cell classes across all datasets.

**Table 1.**
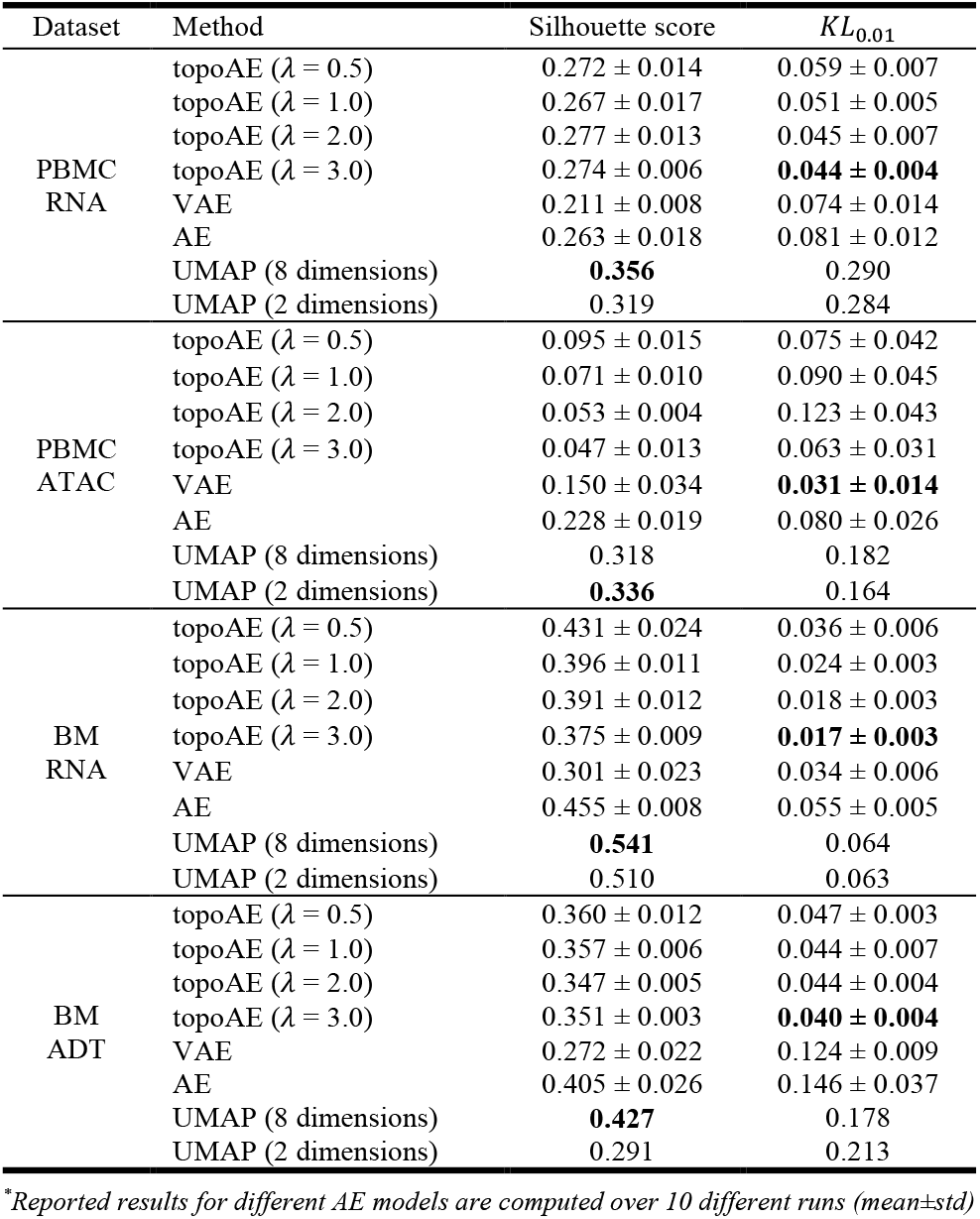
Manifold projection evaluation results

| Dataset | Method | Silhouette score | $KL_{0.01}$ |
| --- | --- | --- | --- |
| PBMC RNA | topoAE ($\lambda = 0.5$) | $0.272 \pm 0.014$ | $0.059 \pm 0.007$ |
| | topoAE ($\lambda = 1.0$) | $0.267 \pm 0.017$ | $0.051 \pm 0.005$ |
| | topoAE ($\lambda = 2.0$) | $0.277 \pm 0.013$ | $0.045 \pm 0.007$ |
| | topoAE ($\lambda = 3.0$) | $0.274 \pm 0.006$ | **$0.044 \pm 0.004$** |
| | VAE | $0.211 \pm 0.008$ | $0.074 \pm 0.014$ |
| | AE | $0.263 \pm 0.018$ | $0.081 \pm 0.012$ |
|  | UMAP (8 dimensions) | **0.356** | 0.290 |
|  | UMAP (2 dimensions) | 0.319 | 0.284 |
| PBMC ATAC | topoAE ($\lambda = 0.5$) | $0.095 \pm 0.015$ | $0.075 \pm 0.042$ |
| | topoAE ($\lambda = 1.0$) | $0.071 \pm 0.010$ | $0.090 \pm 0.045$ |
| | topoAE ($\lambda = 2.0$) | $0.053 \pm 0.004$ | $0.123 \pm 0.043$ |
| | topoAE ($\lambda = 3.0$) | $0.047 \pm 0.013$ | $0.063 \pm 0.031$ |
| | VAE | $0.150 \pm 0.034$ | **$0.031 \pm 0.014$** |
| | AE | $0.228 \pm 0.019$ | $0.080 \pm 0.026$ |
|  | UMAP (8 dimensions) | 0.318 | 0.182 |
|  | UMAP (2 dimensions) | **0.336** | 0.164 |
| BM RNA | topoAE ($\lambda = 0.5$) | $0.431 \pm 0.024$ | $0.036 \pm 0.006$ |
| | topoAE ($\lambda = 1.0$) | $0.396 \pm 0.011$ | $0.024 \pm 0.003$ |
| | topoAE ($\lambda = 2.0$) | $0.391 \pm 0.012$ | $0.018 \pm 0.003$ |
| | topoAE ($\lambda = 3.0$) | $0.375 \pm 0.009$ | **$0.017 \pm 0.003$** |
| | VAE | $0.301 \pm 0.023$ | $0.034 \pm 0.006$ |
| | AE | $0.455 \pm 0.008$ | $0.055 \pm 0.005$ |
|  | UMAP (8 dimensions) | **0.541** | 0.064 |
|  | UMAP (2 dimensions) | 0.510 | 0.063 |
| BM ADT | topoAE ($\lambda = 0.5$) | $0.360 \pm 0.012$ | $0.047 \pm 0.003$ |
| | topoAE ($\lambda = 1.0$) | $0.357 \pm 0.006$ | $0.044 \pm 0.007$ |
| | topoAE ($\lambda = 2.0$) | $0.347 \pm 0.005$ | $0.044 \pm 0.004$ |
| | topoAE ($\lambda = 3.0$) | $0.351 \pm 0.003$ | **$0.040 \pm 0.004$** |
| | VAE | $0.272 \pm 0.022$ | $0.124 \pm 0.009$ |
| | AE | $0.405 \pm 0.026$ | $0.146 \pm 0.037$ |
|  | UMAP (8 dimensions) | **0.427** | 0.178 |
|  | UMAP (2 dimensions) | 0.291 | 0.213 |

\*Reported results for different AE models are computed over 10 different runs (mean ± std)

Further, to qualitatively compare the low-dimensional manifolds produced by each method, we generated two-dimensional UMAP embeddings of the 8-dimensional manifolds of the topoAE, VAE and AE, in comparison with the 2-dimensional UMAP embeddings (Fig. 2). Overall, all methods obtained similar maps with good separation between the cell types, however, VAE could not group the CD8 populations for the PBMC ATAC data. Taken together, topoAE showed better performance in producing low-dimensional manifolds preserving the original density of the data, with comparable performance in terms of cell type separation compared to other autoencoder models.

**Fig. 2.**
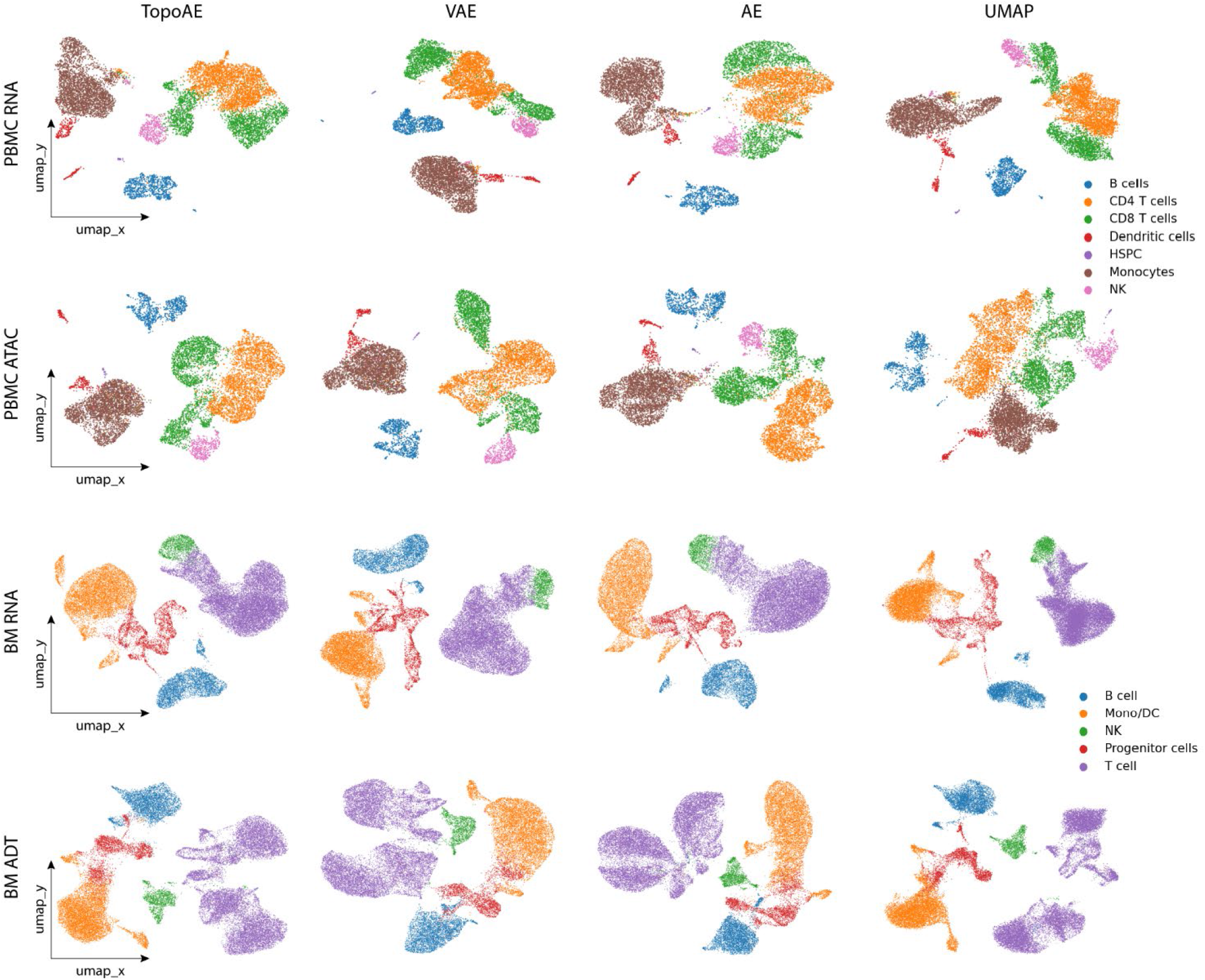
Qualitative comparison of the manifold projection. Plots showing two-dimensional UMAP embeddings of the 8-dimensional manifolds obtained using topoAE, VAE and AE, together with the 2-dimensional embedding obtained using UMAP, for the RNA and ATAC modalities of the **Full PBMC** dataset (top two rows) and the RNA and ADT modalities of the **BM** dataset (bottom two rows). All plots are colored according to the cell classes.

### 3.2 Minimum topological loss ensured manifold alignment instead of superposition

After obtaining the lower-dimensional manifold of each modality using topoAEs, these manifolds are integrated into one common space. We applied the scTopoGAN manifold alignment on the **Full PBMC** dataset, aligning the ATAC modality (source) to the RNA space (target). We observed an inconsistency in the alignment performance when training multiple GANs initialized with different weights. Although their different losses were more or less equal, the Celltype matching score (see Methods) of 40 different GANs was 37.1 ± 16.5% (mean ± standard deviation). We visualized the resulting alignments for the best and the worst models (Fig. 3). Both GANs achieve a good superposition of the ATAC manifold onto the RNA manifold, aligning the ATAC data to match the shape of the RNA data. However, the worst GAN produced a poor alignment of cell classes, e.g. projecting T cells to Monocytes (Fig. 3A-B). Whereas, the best GAN correctly aligns most cell classes (Fig. 3C-D).

**Fig. 3.**
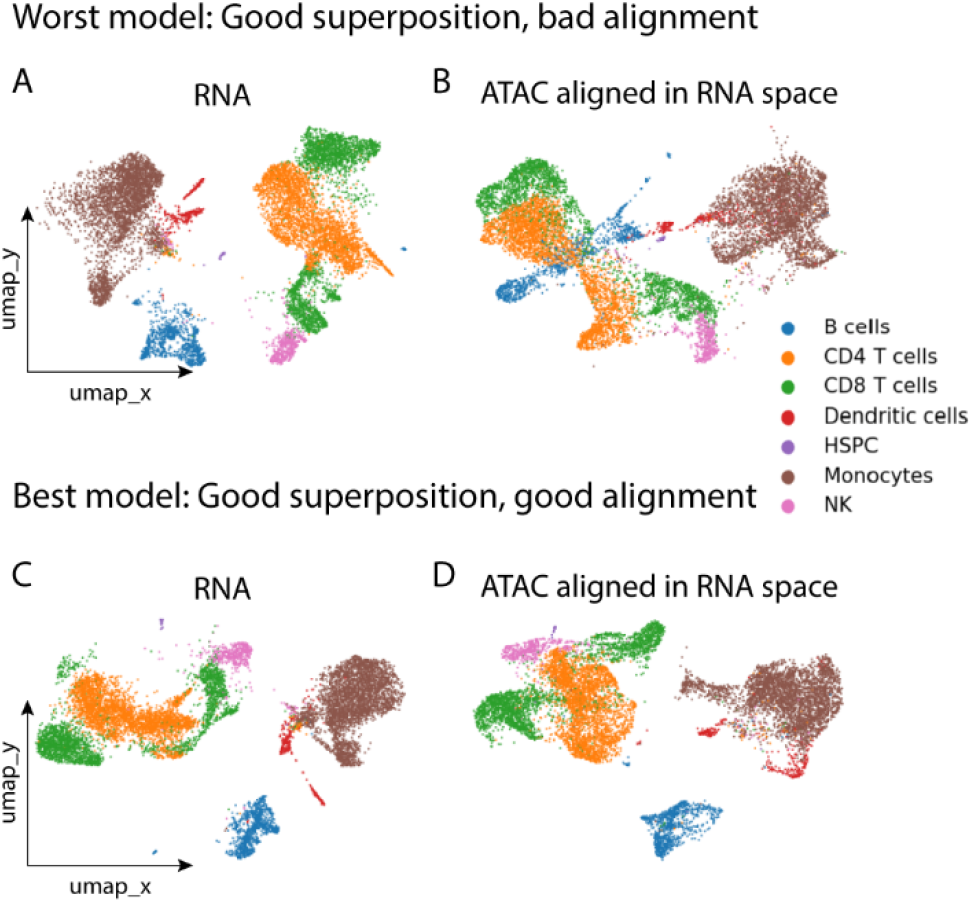
GAN alignment vs superposition. Plots show UMAP embeddings of the final alignment of two GAN models, showing **(A, C)** the RNA modality (target) and **(B, D)** the ATAC modality (source) after being projected and aligned in the RNA space. **(A, B)** Worst GAN model performing good superposition of the two manifolds, but bad alignment in terms of cell classes (8.2% Celltype matching score). **(C, D)** Best GAN model with good alignment projecting the correct cell classes across the two modalities (70.4% Celltype matching score).

To quantify how distorted the source manifold is after projection to the target space, we inspected the topological loss between the source data and the projected source. Using the **Full PBMC** dataset, we performed 40 experiments using identical GAN architectures but different random initializations, trained for 1000 epochs. GAN training showed that the generator and discriminator losses stabilized around 400 epochs. Therefore, we calculated the topological loss from epoch 500 to 1000 every 100 epochs, between the input ATAC (source) manifold and the output ATAC manifold projected onto the RNA space. To interpret the topological losses, we correlated them with the Celltype and the Subcelltype matching scores at the same epochs, resulting in a negative Pearson correlation of -0.73 and -0.67, respectively. This negative correlation indicates that GANs with low topological loss (i.e. preserving the topology of the source data after alignment) tend to produce better manifold alignment. This observation promoted us to train 20 different GANs and select the model with the minimum average topological loss (see Methods) as the final scTopoGAN model. We chose to train 20 base GANs as that showed to cover a wide range of alignment scores.

### 3.3 scTopoGAN outperforms state-of-the-art methods

We benchmarked scTopoGAN against UnionCom and MMD-MA. First, we tested the three methods using all three datasets (**Full PBMC, Partial PBMC** and **BM**), and evaluated the results using the Celltype and Sub-celltype matching scores (Table 2). Shuffling the input data and random sampling of the batches in each iteration (in the case of scTopoGAN) ensured that the cell-cell correspondence is not implicitly captured by any of the methods. In all cases, scTopoGAN outperformed UnionCom and MMD-MA based on both scores, producing joint embeddings with better cell type separation.

**Table 2.** Benchmarking scTopoGAN against UnionCom and MMD-MA

| Dataset | Method | Celltype matching<br>(mean ± std) % | Subcelltype matching<br>(mean ± std) % |
| --- | --- | --- | --- |
| Full<br>PBMC | scTopoGAN | **61.7 ± 8.6** | **41.3 ± 6.5** |
| | UnionCom | 33.8 ± 11.7 | 20.1 ± 10.2 |
| | MMD-MA | 23.3 ± 8.2 | 7.5 ± 7.1 |
| Partial<br>PBMC | scTopoGAN | **72.5 ± 5.1** | **45.1 ± 1.8** |
| | UnionCom | 30.2 ± 7.7 | 15.5 ± 6.4 |
| | MMD-MA | 21.9 ± 9.0 | 6.6 ± 6.9 |
| BM | scTopoGAN | **50.9 ± 14.7** | **22.5 ± 5.4** |
| | UnionCom | 34.7 ± 9.0 | 15.9 ± 6.1 |
| | MMD-MA | 24.9 ± 14.9 | 8.2 ± 7.2 |

*\*Reported results are computed over 10 different runs.*

Further, we qualitatively compared the performance of scTopoGAN, UnionCom and MMD-MA in order to interpret their performances. We visualized the final alignment results for all three datasets (Fig. 4). For the **Full PBMC** and **Partial PBMC** datasets, scTopoGAN showed better mixing of the RNA and ATAC modalities compared to UnionCom and MMD-MA, while keeping the cell classes separable (Fig. 4A-B). The **BM** dataset is more challenging to correctly match the smaller cell classes, however, scTopoGAN produced better alignment and mixing of the RNA and ADT modalities compared to other methods (Fig. 4C).

**Fig. 4.**
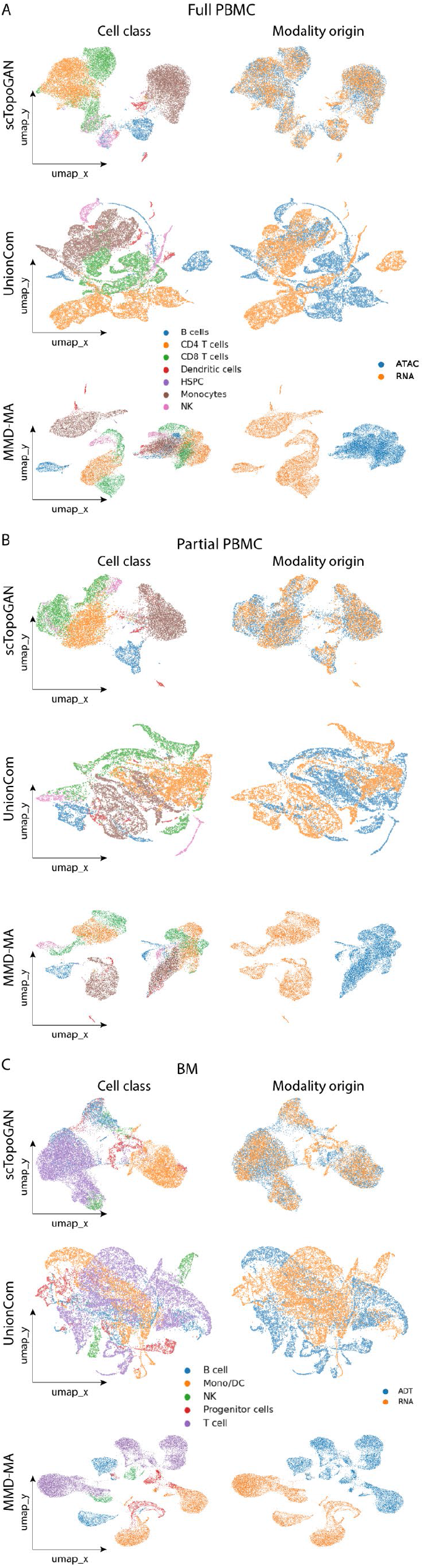
Qualitative comparison of scTopoGAN, UnionCom and MMD-MA. Plots show UMAP embeddings of the final alignment produced by each method when applied on **(A) Full PBMC** dataset, **(B) Partial PBMC** dataset, and **(C) BM** dataset. Each UMAP is plotted twice, once colored with the cell classes showing how well different cell classes are separated, and once colored with the modality of origin showing how well different modalities are mixed.

Finally, we quantified the computational complexity of the different methods. In terms of memory requirements, scTopoGAN memory requirement is almost constant with the number of cells, due to the fixed batch size. For the three dataset (**Full PBMC, Partial PBMC** and **BM**), scTopoGAN is the least demanding method using approximately 2.5 GB for all experiments. MMD-MA used 3.0, 2.6 and 3.0 GB, respectively. While UnionCom required 7.4, 4.6 and 6.4 GB respectively. In terms of computation time, UnionCom was the fastest method with average running times of 1.5, 0.6 and 1.8 hours for the **Full PBMC, Partial PBMC** and **BM** datasets, respectively. scTopoGAN required 1.9, 1.4 and 2.4 hours, respectively. Lastly, MMD-MA running times were 2.1, 1.0 and 2.0 hours, respectively.

## 4 Discussion

We present scTopoGAN, a method to integrate multi-modal single-cell data with non-overlapping cells or features. scTopoGAN is fully unsupervised and relies on the assumption that different single-cell modalities measured from the same tissue have the same underlying manifold, hence the topological structure of these modalities should be similar. To perform manifold alignment, scTopoGAN uses a GAN in combination with a topological loss guiding the selection of the best performing GAN. We would like to stress that although the topological loss idea was inspired based on the evaluation using the cell type annotations, these annotations are not at all used by scTopoGAN (the topological loss is fully unsupervised).

We showed that scTopoGAN outperforms current state-of-the-art methods, UnionCom and MMD-MA. However, the best Celltype matching obtained was ∼70% which shows how difficult and challenging the task of diagonal integration is. Although the topological loss calculation is capable, in most cases, to select the generator model with relatively good alignment performance, it is important to note that training the generator is quite a difficult task since the generator has no ground to link cells between the source and the target data.

For manifold projection, we used topoAEs and showed their ability to preserve the structure of the data in the low-dimensional embedding. topoAEs showed better results compared to VAE, AE and UMAP. However, UMAP still produced the best separation between the cell classes. Nevertheless, this evaluation is biased towards UMAP, as these cell classes were defined using clustering analysis, where UMAP is used to interpret the clusters and annotate them.

Further, it was previously shown that topoAEs have superior performance to PCA and regular autoencoder (Moor *et al*., 2020). Therefore, it might be interesting to explore the applicability of topoAEs in other single cell analysis tasks. One example is trajectory inference studying the differentiation trajectory of cells using scRNA-seq datasets (Saelens *et al*., 2019). Most trajectory inference methods rely on a lower-dimensional representation of the data, where topoAEs can be applied to produce low-dimensional space preserving the topology of the inter-cellular relationships in the data.

The main assumption of scTopoGAN, different modalities measured from the same tissue have the same underlying manifold structure, is not completely true. Although this assumption is based on the fact that the cell pool where different modalities sample from is the same, hence similar cellular structure, different modalities are measuring different molecular features capturing different views of this cellular structure. As a result, the underlying manifolds of each modality are not identical, however, we argue that there is enough similarity between these manifold that can be used to perform the data integration.

A major limitation in the current scTopoGAN workflow is the requirement of training multiple GAN networks in order to choose the best model based on the topological loss. It is evident that the quality of the alignment achieved is limited by the best alignment obtained in this set of GAN models. Here, we trained 20 different GAN models which is computationally expensive and there is no guarantee that the selected GAN model is the best possible solution for the tested dataset. Future improvement in this direction can incorporate the topological loss as a regularization term in the overall loss function of the GAN. This will guide the GAN to minimize the topological loss during training, thus eliminating the need to train multiple GAN models.

In all our experiments, we used the RNA modality as the target modality to which other source modalities (ATAC or ADT) were aligned. The choice of the target modality has an impact on the final alignment performance. Furthermore, we did not fine-tune the hyperparameters used for each dataset. Optimizing these hyperparameters specifically for each dataset may improve the overall results.

In conclusion, scTopoGAN opens new opportunities in studying complex tissues as it represents a step towards better integration of multiple molecular views without the restriction that these are measured from the same cell.

